# Isothermal digital detection of microRNA using background-free molecular circuit

**DOI:** 10.1101/701276

**Authors:** Guillaume Gines, Roberta Menezes, Kaori Nara, Anne-Sophie Kirstetter, Valérie Taly, Yannick Rondelez

## Abstract

MicroRNA, a class of transcripts involved in the regulation of gene expression, are emerging as promising disease-specific biomarkers accessible from tissues or bodily fluids. However, their accurate quantification from biological samples remains challenging. We report a sensitive and quantitative microRNA method using an isothermal amplification chemistry adapted to a droplet digital readout. Building on molecular programming concepts, we design DNA circuit that converts, threshold, amplifies and report the presence of a specific microRNA, down to the femtomolar concentration. Using a leak-absorption mechanism, we were able to suppress non-specific amplification, classically encountered in other exponential amplification reactions. As a result, we demonstrate that this isothermal amplification scheme is adapted to digital counting of microRNA: by partitioning the reaction mixture into water-in-oil droplets, resulting in single microRNA encapsulation and amplification, the method provides absolute target quantification. The modularity of our approach enables to repurpose the assay for various microRNA sequences.

## Introduction

MicroRNA are endogenous short non coding RNA strands involved in post-transcriptional regulation of gene expression (*1*). Discovered 20 years ago, microRNA are emerging as attractive diagnostic, prognostic or predictive biomarkers. Firstly, because accumulating clinical evidence suggests that dysregulation of microRNA is closely related to various diseases including cancers, neuronal or heart diseases, diabetes (*2*) as well as to resistance to chemo or radiotherapy (*3*). Secondly, they are present in bodily fluids, the so-called minimally invasive liquid biopsies (serum, plasma, urine) (*4*). Circulating biomarkers can be assessed readily, potentially enabling a regular follow-up of treatment efficacy, detection of relapses, early diagnostic or large scale population screening (*5*). Finally, these nucleic acids with known sequences are amenable to rational and rapid design of generic detection tests.

The detection of circulating microRNA in blood is still a challenging task as they are highly diluted. Besides, diseased-related microRNA must be accurately quantified to leverage their use in clinical procedures. Hence, the precise measurement of microRNA concentrations remains the critical bottleneck, encouraging the development of sensitive, specific and quantitative detection technologies. The current gold standard for sensitive microRNA detection is the reverse transcription-quantitative polymerase chain reaction (RT-qPCR) (*6*). Despite the high sensitivity of RT-qPCR, the technique possesses inescapable drawbacks: i) the RT step is known to introduce significant bias in the quantification (*7*–*9*); ii) primers and probe must be designed for each target and rely on sophisticated design due to the short length of the target; iii) the thermocycling protocol should be optimized for each assay; iv) the target itself is amplified and is therefore a hazardous source of carry-over contamination (*10*); v) PCR reaction is known to be inhibited by biological samples (*11*). Additionally, the procedure relies on a real-time tracking of the amplification and requires standard calibration, thus giving only access to a relative estimate of the amount of microRNA.

This last issue can in principle be addressed with a digital quantification (such as droplet-PCR (*12*)) based on the isolation of single nucleic acid molecules in microcompartments. Although dPCR emerges as a powerful method for precise and absolute quantification of nucleic acids, it retains the weaknesses of RT-qPCR for microRNA assessment (*13*). Isothermal alternatives (*14*) have been proposed (EXPAR (*15, 16*), LAMP (*17*), RCA (*18*), HCR (*19*), etc.) that rely on simpler one-step protocols and free themselves from the reverse transcription reaction. Among these isothermal strategies, EXPAR (EXPonential Amplification Reaction) has gained attention due to its simplicity and competing sensitivity. However, the exponential nature of the reaction renders it prone to leaky reactions, eventually triggering the production of amplified products from spurious reactions (*20, 21*). These nonspecific amplifications are always observed with a short delay after the target-triggered reaction. This issue makes such amplification chemistry ill-adapted to a digital readout where only the target-containing compartments are expected to produce a positive signal (Supplementary Figure 1). Preheating separately the substrates (templates and dNTP) and the enzymes (polymerase, endonuclease) may delay target-independent amplification, however a digital readout would still require a very precise control over the incubation time to limit the occurrence of false positive compartments (*16*). Hairpin-shaped templates have been shown to improve the specificity over homologous sequences, although target-independent amplification is not delayed (*22*). Several additives such as graphene oxide (*23*), tetramethylammonium chloride (*24*) or single strand binding protein (*24, 25*) have been proposed to decrease these spurious reactions and increase assay sensitivity. Recently, Urtel et al. successfully suppressed background amplification using a template lacking one of the four nucleotide (dA-free template), a three-letter nicking site and dATP-free amplification mixture (*26*). Similarly, locked nucleic acid and peptide nucleic acid modification of the 3’ domain of EXPAR template have been used to reduce background amplification (*25, 27*). Nevertheless, no biosensing application has been validated yet with these last two approaches (*25*–*27*).

Besides enzyme-based amplification reaction, enzyme-free system have been reported using toehold-mediated strand displacement reactions. These reactions are also prone to leak, coming from the spurious release of output strands, which is detrimental for the sensitivity in sensing applications (*28*). Many strategies have been reported to mitigate the leaks, encompassing clamped (*29*–*31*) or long (*32*) domains, the introduction of mismatches (*33*), or using ultrapure DNA complexes (*28, 31, 32*). However, in comparison with the enzyme-powered approaches, most enzyme-free circuits display limited sensitivity as a consequence of lower signal amplification efficiency. Herein, we propose a background-free isothermal amplification technique relying on a DNA-based molecular program specifically designed to catalyze a target-triggered signal amplification, while avoiding nonspecific amplification reactions. The amplification system is coupled to a catalytic degradation mechanism that continuously absorbs products stemming from leaky reaction. Applied to the detection of microRNA, we demonstrate the high sensitivity of this amplification chemistry down to femtomolar concentrations. The suppression of nonspecific amplification reaction allows the robust droplet digital detection of microRNA targets and their absolute end-point quantification.

## Results

### Molecular program designed for microRNA detection

Molecular programming is an emerging discipline that involves the design of artificial biomolecular circuits performing information-processing tasks. It uses the predictable Watson and Crick base-pairing of synthetic DNA oligonucleotides to enable the rational design of molecular circuits (*34, 35*). We use here a versatile molecular programming language named PEN-DNA toolbox (Polymerase Exonuclease Nickase-Dynamic Network Assembly) (*36, 37*). It employs a set of short oligonucleotides (templates) encoding the topology of the network and a mixture of enzymes that catalyze the production and degradation of activator/inhibitor strands. Using this set of reaction modules, we designed a generic molecular program dedicated to the detection of microRNA.

Figure 1 presents the connectivity of the circuit (cf. also Supplementary Figure 2 for a more detailed mechanism): the universal signal amplification system is composed of two templates: an autocatalytic template (aT) and a pseudotemplate (pT) (*38*). The aT is made of a dual-repeat sequence catalyzing the exponential replication of a 12-mer oligonucleotide (signal strands, s-strand) via a polymerization/nicking cycles. It is well-known that such exponential reaction are prone to non-specific amplification, meaning that even in absence of the input signal strand, the autocatalysis will eventually occur due to spurious reactions, also known as leak (*20*). The pT catalyzes the deactivation of the s-strand produced from nonspecific reactions and is therefore essential to avoid unwanted amplification (*38*). The molecular system aT:pT acts as a bistable switch (Supplementary Figure 3): in absence of the s-strand input, the leak is absorbed by the pT and amplification is not observed. If the concentration of s-strand exceeds a certain threshold (tuned by the ratio aT/pT), the pT-mediated degradation path is saturated and amplification occurs. In order to make this bistable switch responsive to the microRNA target, a converter template (cT) is connected upstream to the aT: upon hybridization of the microRNA to the input part of the cT, the latter linearly produces trigger strands (by polymerization/nicking reactions), which in turn bind and activate the autocatalytic reaction on the aT. Downstream to the aT, a reporting template (rT) captures the amplified s-strands to produce a fluorescence signal. The dynamical nature of the molecular circuit is preserved by the presence of an exonuclease that degrades all produced strands, keeping the system out of equilibrium.

**Figure 1.**
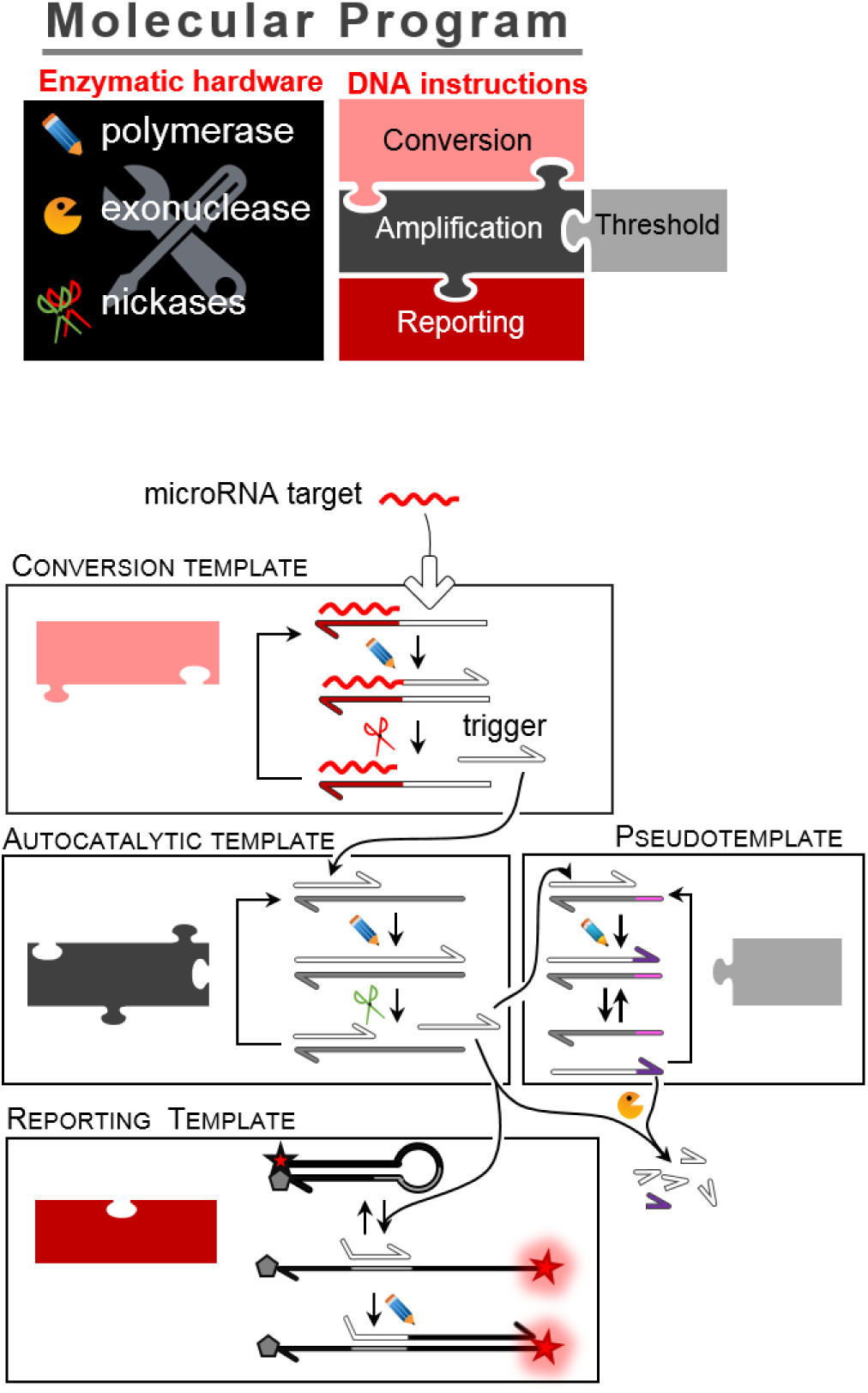
A molecular program dedicated to the detection of microRNA. A 4-template DNA circuit encodes the connectivity of the molecular program, whose reactions are catalyzed by a set of enzymes (polymerase, exonuclease, endonucleases): the cT converts the target microRNA to a generic s-strand sequence; the aT exponentially amplifies the s-strand sequence; to avoid target-independent amplification, the pT drives the deactivation of the small amount of triggers originating from leaky reactions; the rT translates the s-strand sequences into a fluorescence signal.

### Background-free detection of the microRNA Let-7a

Figure *2* evaluates the sensitivity of this approach for the real-time detection of Let-7a. After optimizing the experimental conditions (Supplementary Figure 4), the four templates were mixed together with the enzymatic processor and spiked with a concentration of synthetic RNA target Let-7a ranging from 0 to 1 nM (Figure *2*A). The fluorescence of the rT is monitored in real-time with a PCR thermocycler set at a constant temperature of 50°C. The negative control (no target) does not produce a positive signal more than 20 hours (the experiment duration).

**Figure 2.**
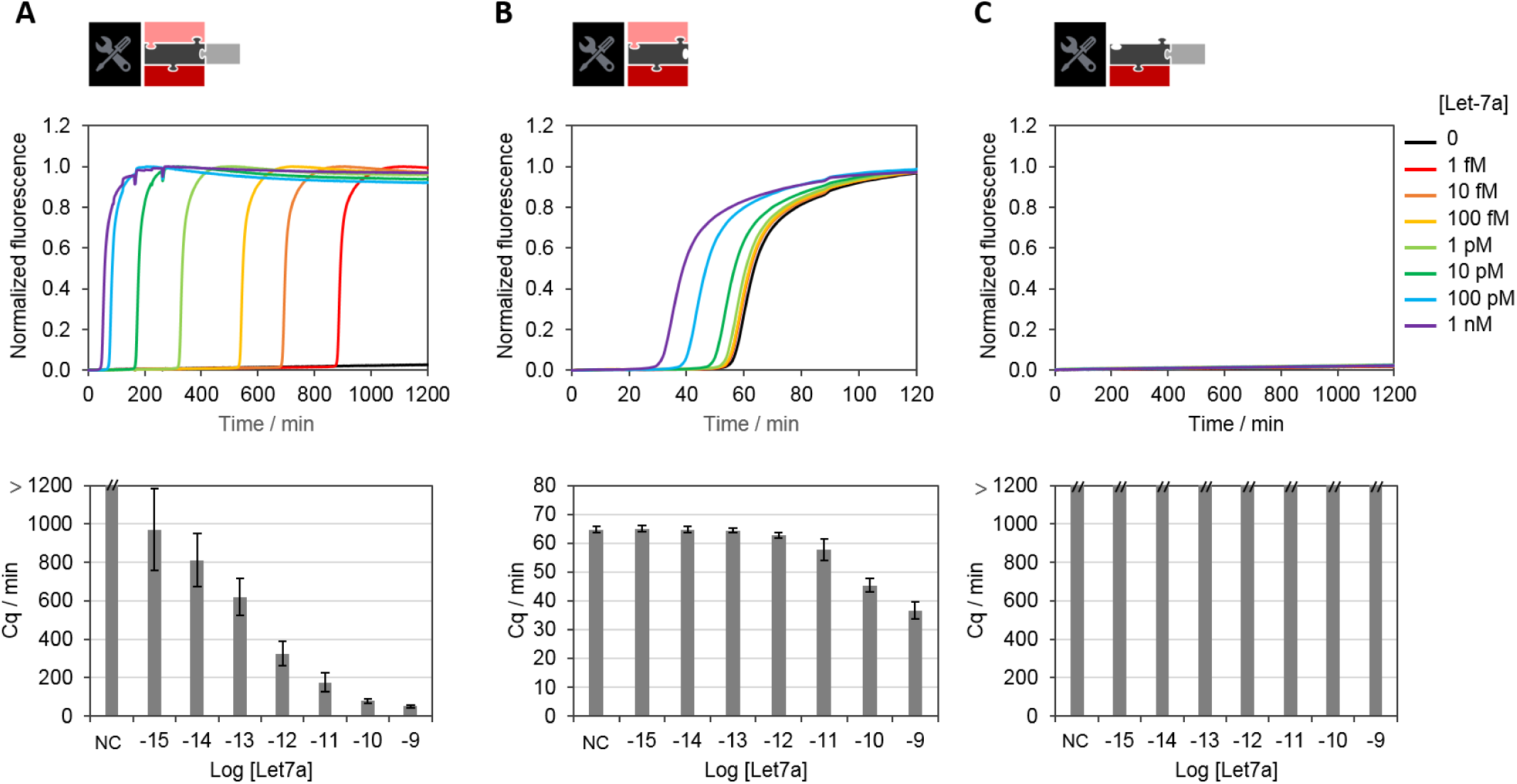
In-solution detection of the microRNA Let7a. (**A**) Detection with the 4-template molecular program, (**B**) the molecular program without the pseudotemplate or (C) without the converter template. The amplification reaction is monitored in real-time and the amplification time (Cq) is plotted as a function of the Let-7a concentration.

The limit of detection of the assay is around 1 fM and the dynamic range spans 6 orders of magnitude from 1 fM to 100 pM. The absence of amplification when the cT is missing (Figure *2*C) validates the specificity of the target-triggered amplification. In absence of the pT (Figure *2*B), the sensitivity is negatively affected with a limit of detection of 3.7 pM (estimated from three standard deviation from the mean amplification time of the negative control). These results demonstrate the importance of this active leak-absorption mechanism for controlling the amplification threshold and thus eliminating background amplification. However, the active consumption of s-strands increases considerably the time of the assay. For instance, it takes 16 hours to trigger the amplification reaction from 1 fM of Let7a target, which is too slow for most routine procedures.

### Droplet digital detection for absolute microRNA quantification

To circumvent this issue, we converted the analog readout, based on relative measurements, to a digital readout where single molecules are detected after being isolated in independent compartments (Figure *3*) (*12, 39*). Using a flow focusing microfluidic device, the master mix spiked with a known concentration of Let-7a was partitioned in picoliter-sized water-in-oil droplets (Figure *3*A). The resulting monodisperse emulsion was incubated at 50°C for 200 min, after which the droplets were imaged by fluorescence microscopy (Figure *3*B) and the concentration of target computed from the Poisson statistics (Supplementary Figure 5). We observe a linear correlation between the spiked microRNA concentration and the concentration computed according to the Poisson law (Figure *3*D and Supplementary Figure 6). It is important to note that another benefit of the partitioning is that the time necessary to amplify the signal from a single molecule depends only on the volume of the compartments, not on the concentration in the initial sample. For droplets of 0.5 pL (∼10 µm in diameter), the concentration of a single molecule corresponds to 3 pM, thus decreasing the incubation time down to ∼3 hours (Figure *2*A). The modularity of the DNA circuit enables us to repurpose the molecular program by redesigning solely the cT to the miRNA target of choice (e.g. miR-92a or cel-miR-39 in Figure *3*E), while the other reaction components (aT, pT, rT) remain unchanged.

**Figure 3.**
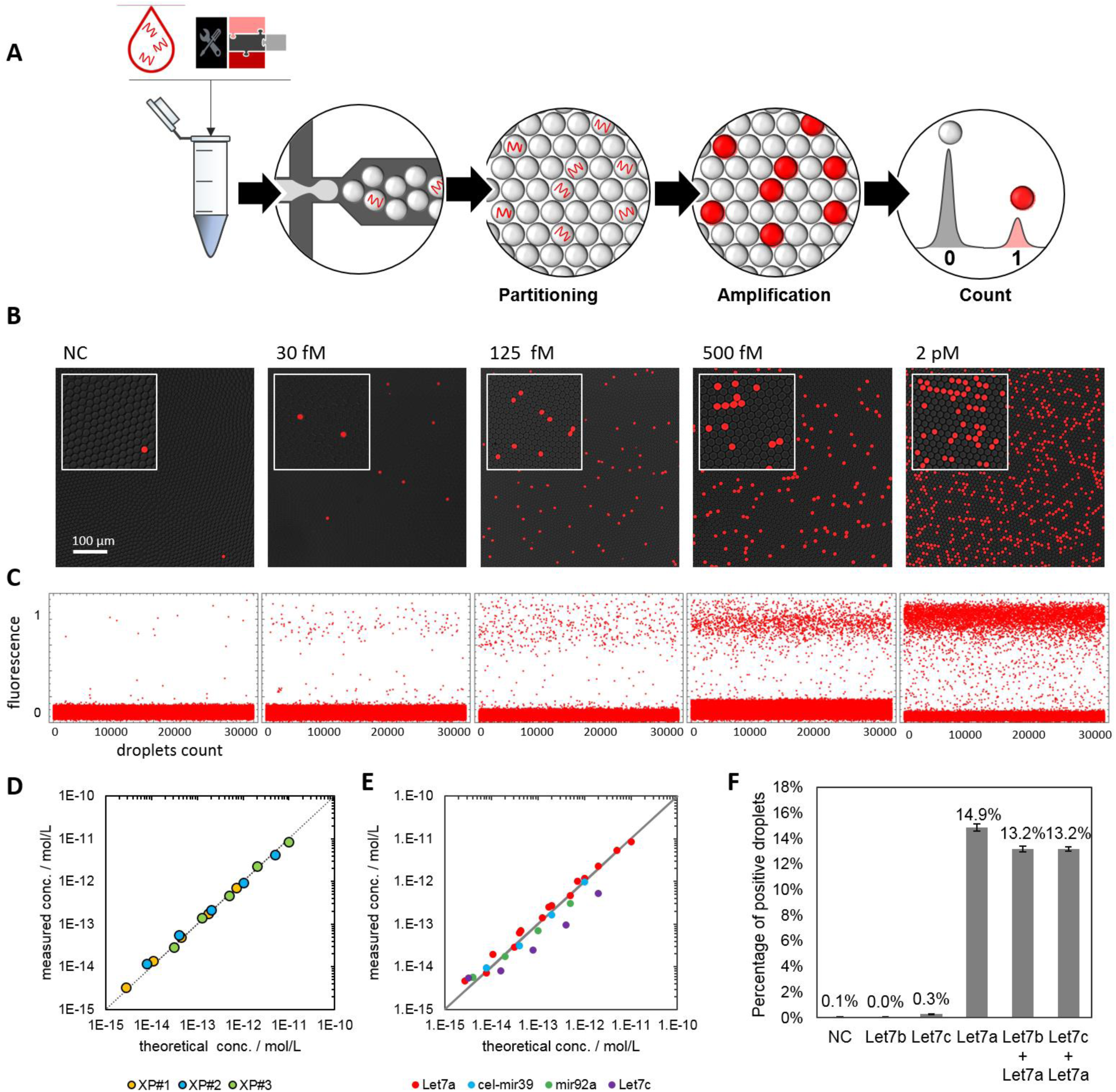
Digital detection of microRNA. (**A**) The sample is mixed with the molecular program and partitioned into monodisperse droplets resulting in the random distribution of the microRNA targets among the compartments. After incubation, the droplets are sandwiched between two glass slides and imaged by fluorescence microscopy. The droplets having received at least one target exhibit a positive fluorescence signal (1), while the others remain negative (0), allowing absolute concentration calculation using Poisson statistics. (**B**) Fluorescence snapshot of emulsified samples spiked with increasing concentrations of Let-7a after amplification. (**C**) Analysis of 30000 droplets. (**D**) Plot of the linear relationship between the expected (theoretically spiked) and the experimentally measured target concentration. The data were obtained from three independent experiments “XP” and the measured concentration corresponds to the concentration calculated from the Poisson statistics (cf. Supplementary Figure 5) subtracted by the limit of the blank (cf. Supplementary Figure 6) (**E**) By adapting the converter template, the assay is repurposable for other microRNA. (**F**) The assay specificity is evaluated from the cross reactivity of Let-7a over Let-7c (single mismatch) and Let-7b (2 mismatches) and in mixtures (1 pM each). Error bars represent the 95 % confidence interval over digital quantification.

A major concern related to the quantification of microRNA is the high sequence homology between microRNA sequences. We thus evaluated the specificity of the current detection method over the Let-7 family. Figure *3*F shows a clear discrimination between the Let-7a sequence and analog sequences containing a single (in the case of Let-7c) or two (in the case of Let-7b) mismatched bases. The selectivity of the current method, estimated to be superior to 97 %, is among the best ratios described in the literature for microRNA digital assays (*40*–*42*).

The high selectivity can be explained by the dynamics of the systems that needs to produce a concentration of trigger exceeding a given threshold to initiate the amplification. While the production speed of s-strand is maximum for the full matching duplex cT/microRNA, it is considerably reduced with single-base modified targets (Supplementary Figure 7). The dynamical threshold set by the pseudotemplate deactivation mechanism acts as an enhancer of the specificity because only activated cT producing triggers at a high rate will eventually initiate the amplification.

### Background-free amplification ensures robust digital end-point analysis

As digital assays rely on an end-point analysis, it is crucial to have a time window large enough to discriminate the target-containing droplets (exhibiting a positive signal) from the target-free droplets. As mentioned above, most isothermal nucleic acid amplification techniques cannot be transposed to a digital format due to nonspecific reactions that eventually trigger the amplification in all microcompartments, irrespective of the presence of the target (*20, 40*). Figure 4 shows that in absence of pT, the droplets turn ON in less than an hour, independently of the presence of the target, the frequency of false positive droplets (red curve) reaching almost 50 %. This result is consistent with a previously described EXPAR system (*40*). Between 10 and 13 nM of pT, we observe a significant decrease of the false positive fraction: the pT absorbs part of the leak, but the residual leak is strong enough to unspecifically trigger the amplification in some droplets, resulting in an increase over time of the false positive ratio. By setting the pT concentration above 16 nM and thus raising the amplification threshold, the self-start was completely suppressed. It is important to note that from 16 to 22 nM of pT, the quantification of Let7a concentration is not affected (green curves). This suggests that the digital assay is tolerant to small variations of the molecular program composition. Thanks to this amplification threshold, which avoids unwanted amplification, the present method is robust with respect to incubation time with a time window spanning several hours.

**Figure 4.**
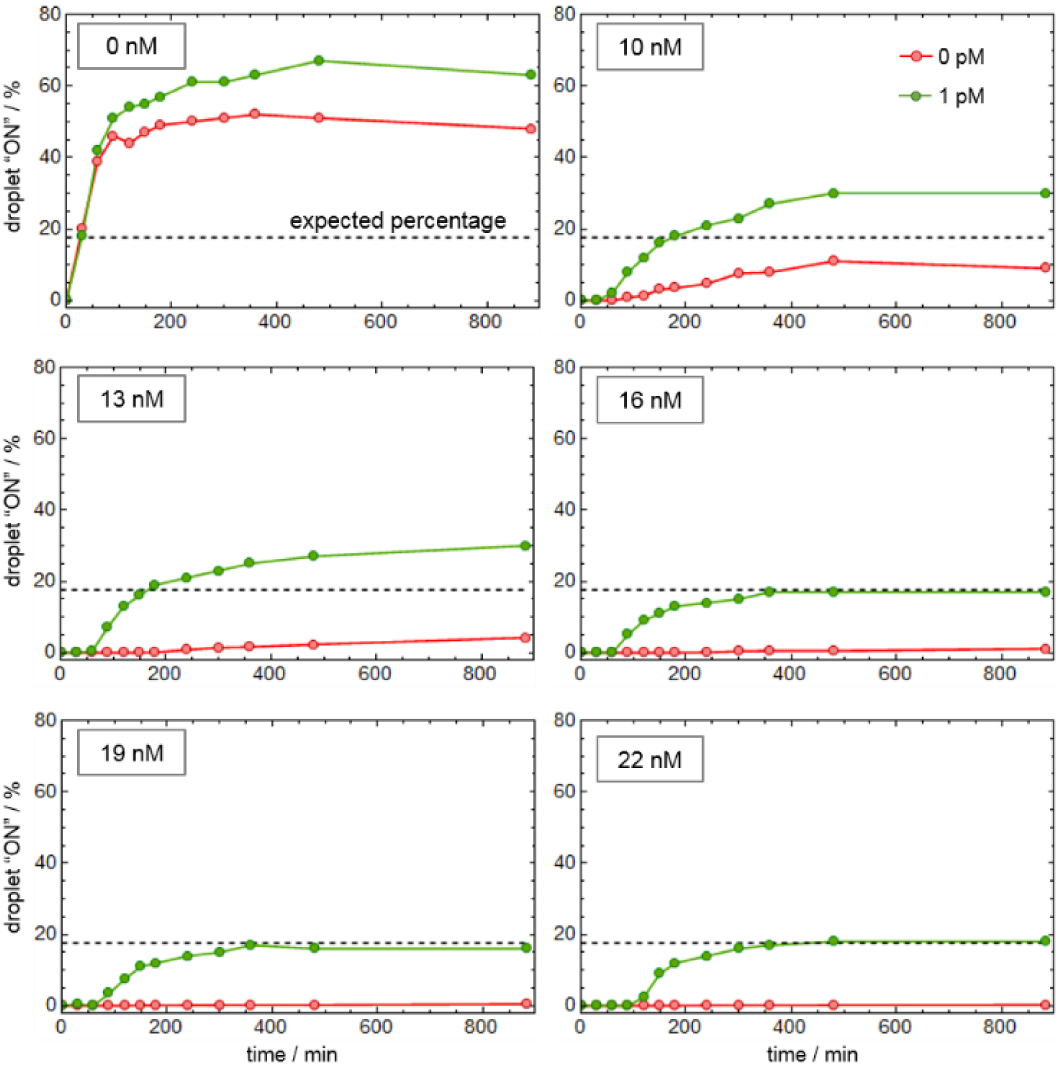
The elimination of non-specific amplification reaction allows robust digital detection of microRNA targets. The molecular program with 0 to 22 nM of pT is spiked with 0 or 1 pM of synthetic Let-7a before partitioning. The droplets are incubated at 50 °C and imaged by fluorescent at different time point to extract the fraction of positive droplets. The dashed line represents the expected percentage given a target concentration of 1 pM and a droplet diameter of ∼ 9 µm.

### Test and validation with total RNA extracts

We then evaluated the possibility to detect endogenous microRNA from human cells. We quantified the microRNA Let-7a from human colon total RNA using the isothermal digital assay. Figure 5a shows a linear relationship between the total RNA concentration (ranging from 0 to 8 ng/µL) and the measured Let-7a concentration (R^2^ > 0.99). Similarly, the quantification of Let-7a using RT-qPCR is linearly correlated to the initial extract concentration. Although the quantification of the Let7a microRNA is consistent between both techniques, the measured concentration is significantly higher with the qPCR assay. The quantification by qPCR is estimated from the amplification time (Cq), which is compared to the ones of standard concentrations (Supplementary Figure 8), assuming the amplification efficiency is the same for both the samples and the standards. However, it has been previously highlighted that imprecision in the PCR calibration or modification of the sample composition significantly affect the quantification (*43*). This is arguably the origin of the overestimation of the concentration of Let-7a observed here using RT-qPCR. We calculated the precision of our method to be > 76 % (56 % for RT-qPCR, calculated from the mean value over the three extract concentration tested, Figure 5b). This is in accordance with previous studies showing the improved accuracy of digital PCR in comparison to qPCR (*42*–*44*). As a negative control experiment, the digital quantification of the non-human microRNA mir-39ce using the cognate DNA circuit (presented in Figure *3*e), shows the absence of this target in the human extract. Overall, this demonstrates the accuracy of the assay in quantifying endogenous microRNA.

**Figure 5.**
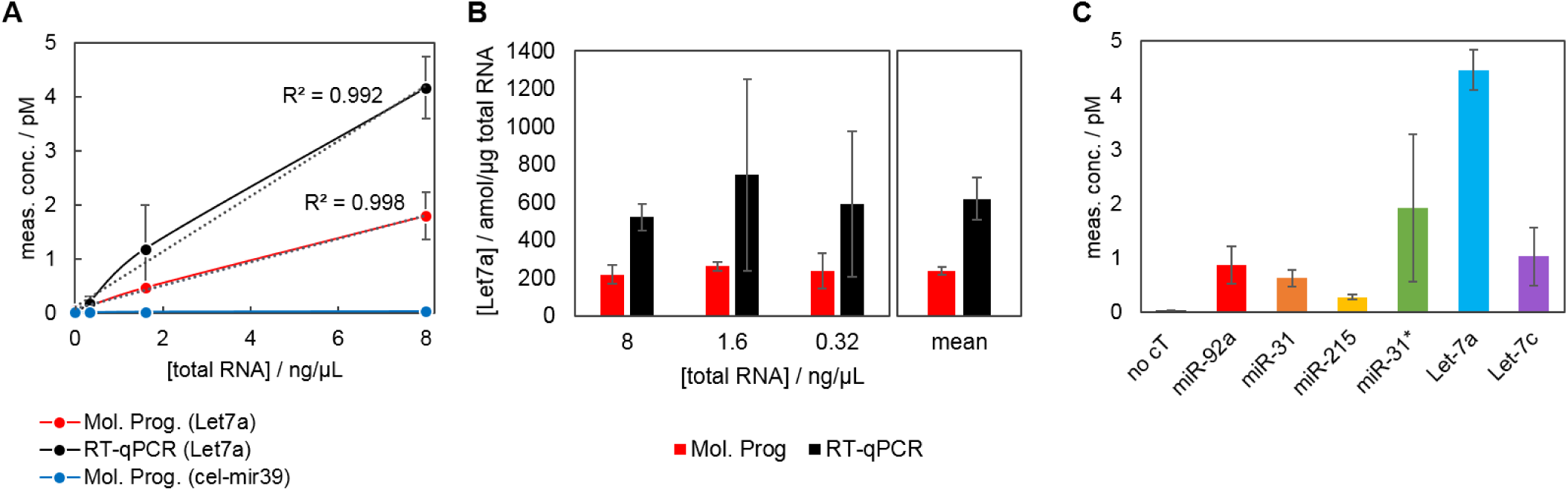
microRNA detection from human colon total RNA extract. (**A**) Comparative quantification of microRNA (Let-7a or mir-39ce) using the proposed MP-based digital assay and RT-qPCR (the error bar are calculated from three independent experiments. (**B**) The Let-7a concentration (in amol/ng of total RNA) is calculated for each total RNA concentration, allowing to determine the precision of the quantification (corresponding to the standard deviation over three replicates of three extract concentrations. (**C**) Multiple microRNA are quantified from 10 ng/µL of total RNA extract.

## Discussion

In conclusion, we present a molecular program-based isothermal amplification strategy that is able to abolish background amplification, thereby enabling digital quantification. The present assay provides sensitive, specific and quantitative measurement of microRNA. Based on a one-step procedure, it reduces sample manipulations and therefore the risk of carry-over contamination. Contamination issues are further reduced by the fact that the system relies on a signal amplification mechanism rather than on the direct replication of the target sequence, as in PCR-based approaches. Our versatile DNA-based circuit can be theoretically repurposed for any microRNA of interest through the simple re-design of a single oligonucleotide. All other circuit components are identical for all targeted microRNA, thus eliminating the need for primers and probe design and reducing the assay cost and optimization. It is to note that the leak absorption mechanism makes the amplification trigger slower compared to some reported EXPAR approaches. In the future, it should be possible to combine it with other leak-reducing amplification strategies (using for instance modified templates(*27*), specific nickases (*26*) or by engineering the DNA polymerase to be less leaky (*45*)). This would decrease the required concentration of pseudotemplate needed to absorb the leak and thus speed up the amplification. Nevertheless, the digital format already significantly decreases the assay time. Digitalization also de-couple the assay duration from the target concentration, with a typical time down to 3 hours in the current implementation.

Because of its sensitivity and quantitativity, we anticipate that this digital isothermal amplification method may be a promising tool in academic and clinical research.

## Materials and Methods

### Chemicals

Oligonucleotides (templates and synthetic microRNA) were purchased from Biomers (Germany). The sequences were purified by HPLC and checked by matrix-assisted laser desorption/ionization mass spectrometry. Templates were designed according to the rules previously described [23], [25]. All template sequences (aT, pT, rT) are protected from the degradation by the exonuclease by 5’ phosphorothioate backbone modifications. A 3’ blocking moiety (phosphate group for aT, pT, cT and quencher for rT) is used to avoid nonspecific polymerization. The Supplementary Table 1 recapitulates all the sequences used throughout the study. Nb.BsmI and Nt.BstNBI nicking enzymes, Vent(exo-) DNA polymerase and BSA were purchased from New England Biolabs (NEB). A 10-fold dilution of Nt.BstNBI was prepared by dissolving the stock enzyme in Diluent A (NEB) supplemented with 0.1 % Triton X-100. The exonuclease ttRecJ was expressed and purified by chromatography according to a previously published protocol (*46*). The enzyme is stored at 1.53 µM in Diluent A + 0.1% Triton X-100. All the proteins were stored at −20°C. Human colon total RNA (Thermofisher Scientific) was aliquoted at 13 µg/mL and stored at −20°C

### RT-qPCR procedure

Quantification was performed by real-time PCR using the Bio-Rad C1000 Touch™ Thermocycler and CFX96™ Real-Time System. A 2-step RT-qPCR protocol was performed according to Life Technologies™ Taqman^®^ MicroRNA Assays instructions and utilizing primers for let-7a (assay ID 000377). The reverse transcription program consists of: 16°C for 30 min, 42°C for 30 min and 85°C for 5 min. The PCR cycling program consists of 1× at 95°C for 10 min, and 60× cycles of 15 s at 95°C and 1 min at 60°C, for denaturing/annealing plus extensions. Default threshold settings were used as threshold cycle (Ct) data. The Ct is the fractional cycle number at which the fluorescence passes the fixed threshold. The Ct is presented as mean±SD, for 3 replicate experiments.

### Reaction mixture assembly

All reaction mixtures were assembled at 4°C in 200 µL PCR tubes: the templates are mixed with the reaction buffer (20 mM Tris HCl pH 7.9, 10 mM (NH4)2SO4, 40 mM KCl, 10 mM MgSO4, 50 µM each dNTP, 0.1% (w/v) Synperonic F 104, 2 µM netropsin, all purchased from Sigma Aldrich) and the BSA (200 µg/mL). After homogenization, the enzymes are added (300 u/mL Nb.BsmI, 10 u/mL Nt.BstNBI, 80 u/mL Vent(exo-), 23 nM ttRecJ). Each sample was spiked with the microRNA target, serially diluted in 1X Tris-EDTA buffer (Sigma Aldrich) using low-binding DNA tips (Eppendorf). The samples (bulk or emulsion) were incubated at 50°C in a qPCR thermocycler (CFX96 Touch, Bio-Rad) and the fluorescence was recorded in real-time. For bulk experiments, the time traces were normalized and the Cq (amplification starting times) determined as 10 % of the maximum fluorescence signal.

### Droplets generation

A 2-inlet (one for the oil, one for the aqueous sample) flow-focusing microfluidic mold was prepared with standard soft lithography techniques using SU-8 photoresist (MicroChem Corp., MA, USA) patterned on a 4-inch silicon wafer. A 10:1 mixture of Sylgard 184 PDMS resin (40 g)/crosslinker (4 g) (Dow Corning, MI, USA) is poured on the mold, degassed under vacuum and baked for 2 hours at 70°C. After curing, the PDMS was peeled off from the wafer and the inlets and outlet holes of 1.5 mm diameter were punched with a biopsy punch (Integra Miltex, PA, USA). The PDMS layer was bound onto a 1 mm thick glass slide (Paul Marienfeld GmbH & Co. K.G., Germany) immediately after oxygen plasma treatment. Finally, the chip underwent a second baking at 200 °C for 5 hours to make the channels hydrophobic [32]. The microfluidic chip details are presented in Supplementary Figure 9. The aqueous sample phase and the continuous phase composed of fluorinated oil (Novec-7500, 3M) containing 1% (w/w) fluorosurfactant (RAN Biotechnologies, MA, USA) were mixed on chip using a pressure pump controller MFCS-EZ (Fluigent, France) and 200 µm diameter PTFE tubing (C.I.L., France) to generate 0.5 pL droplets by hydrodynamic flow focusing.

### Droplets imaging

After incubation, the droplets were imaged by fluorescence microscopy. The bottom slide (76×52×1 mm glass slide) was spin-coated with 200 µL Cytop CTL-809M (Asahi Glass) and dried at 180 °C for 2 hours. The emulsion was sandwiched with a 0.17 mm thick coverslip treated with Aquapel. 10 µm polystyrene particles (Polysciences, Inc., PA, USA) were used as spacer to sustain the top slide and avoid the compression of the emulsion. The imaging chamber was finally sealed with an epoxy glue (Sader) and images were acquired using an epifluorescence microscope Nikon Eclipse Ti equipped with a motorized XY stage (Nikon), a camera Nikon DS-Qi2, an apochromatic 10X (N.A. 0.45) (Nikon) and a CoolLed pE-4000 illumination source. Composite images were generated with the open source ImageJ software. Quantitative data were extracted from the microscopy images using the Mathematica software (Wolfram), following the procedure detailed in the Supplementary Figure 5. The average number of microRNA target per droplet λ is calculated from the frequency of positive droplets (F_pos_) using the Poisson law:

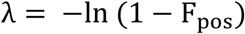

This allows the computation of the microRNA concentration:

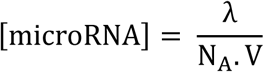

where N_A_ is the Avogadro number and V the volume of the droplets (in L). The 95% confidence interval is given by the uncertainty on a binomial proportion:

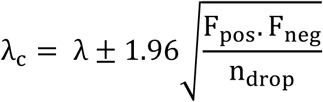

where F_pos_ and F_neg_ correspond to the frequency of positive and negative droplets respectively and n_drop_ is the total number of droplets analyzed.

## Acknowledgments

The authors wish to thank the reviewers of the 25^th^ International Conference on DNA Computing and Molecular Programming for their insight and advices to improve the quality of this manuscript.

## Author contributions

G.G. and R.M. designed the study, performed experiments and analyzed the data. K.N. and A.-S.K. carried out the optimization of experimental conditions. V.T. provided expertise on droplet microfluidics and microRNA markers and supervised this work. Y.R. conceived, designed and supervised the study. G.G. and Y.R. wrote the manuscript. All authors discussed the results and commented on the manuscript.

## Supplementary Materials

Supplementary Figure 1: Basic EXPAR system with hot start.

Supplementary Figure 2. Detailed chemical reaction network of the molecular program for the detection of microRNA.

Supplementary Figure 3. Bistable amplification switch.

Supplementary Figure 4. Experimental conditions optimizations.

Supplementary Figure 5: Droplets analysis.

Supplementary Figure 6: Extended data from the Figure 3d.

Supplementary Figure 7. Specificity of the trigger production.

Supplementary Figure 8. RT-qPCR calibration.

Supplementary Figure 9. Microfluidic chip design.

Supplementary Table 1. Oligonucleotides sequences used throughout the study.

