## Supplementary Material for "Isothermal digital detection of microRNA using background-free molecular circuit"

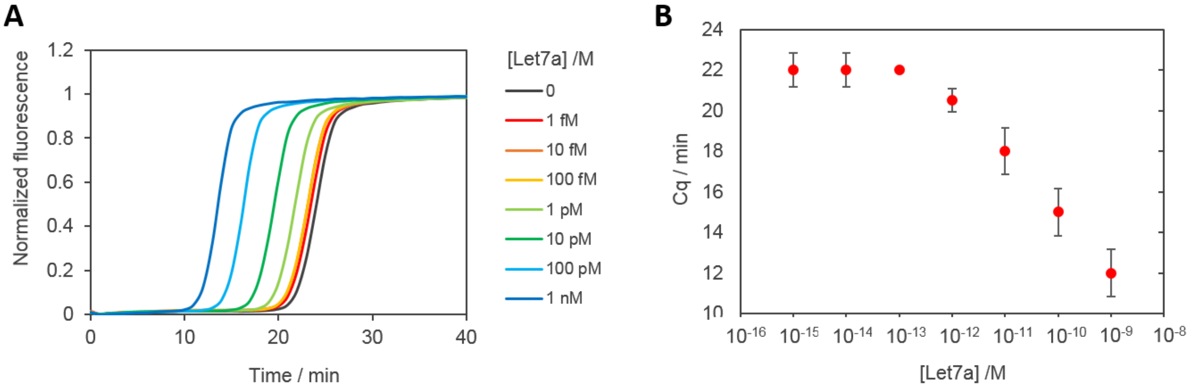


**Supplementary Figure 1: Basic EXPAR system with hot start.** We reproduced the experiment from Zhang et al. using the exponential amplification reaction (EXPAR) based on a dual-repeat template, which enables the replication of the Let-7a sequence using extension (Vent(exo-) DNA polymerase and nicking (Nt.BstNBI nicking enzyme) cycles. Briefly, the reaction mixture was prepared separating the DNA template and dNTP from the enzymes, which has been shown by the authors to delay the self-amplification reaction (in absence of target). Part A. consisted in 1X NEBuffer 3.1, 200 nM EXPAR template (Supplementary Table **1**), 500 µM dNTP (NEB), 1.6 u/µL RNAse inhibitor murine (NEB) and the synthetic Let-7a RNA target at various concentrations. The part B contained 2X ThermoPol buffer (NEB), 800 u/mL Nt.BstNBI, 100 u/mL Vent(exo-) DNA polymerase and 2X EvaGreen (Biotium). Samples are prepared by pre-incubating separately the two mixtures at 55°C for 10 min before combining then (5 µL each), following by a quick vortex mixing. The EXPAR reaction is monitored in real-time at 55°C. (**A**) Amplification curves obtained for various concentrations of Let-7a. (**B**) The Cq (corresponding to the time where the curves reach 10 % of the maximum fluorescence intensity) are plotted as a function of the initial target concentration (the error bars correspond to the standard deviation from three independent experiments, each in duplicate).

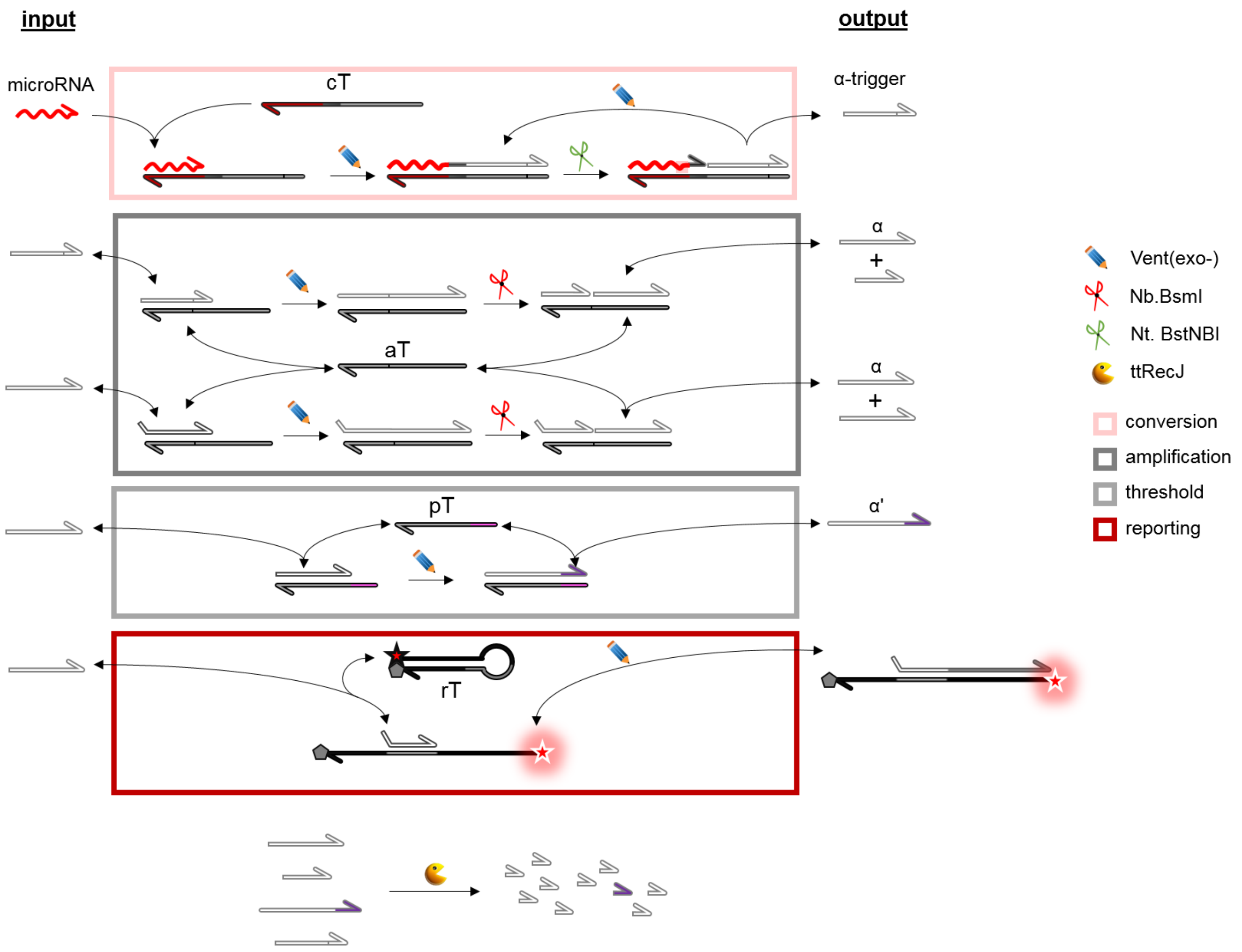


**Supplementary Figure 2. Detailed chemical reaction network of the molecular program for the detection of microRNA.** The molecular program is composed of a set of 4 templates encoding production and degradation of short DNA strands, these reactions being catalyzed by mixture of enzymes. The conversion template (cT) specific of a microRNA sequence, hybridizes to the corresponding target on its input part (3'). The RNA sequence is used as a primer and is extended by the polymerase along the cT. Upon nicking by the enzyme Nt.BstNBI, an α-trigger is released. The catalytic elongation/nicking cycles induce the linear production this trigger. Upon binding of the trigger to the 3' part of the autocatalytic template (aT), the elongation by the polymerase and nicking of the duplex by the enzyme Nb.BsmI produce a molecule of α, which is therefore exponentially amplified. To avoid this autocatalytic reaction to self-start from spurious reactions, a pseudotemplate (pT) is introduced. The pT catalyzes the deactivation of α strands by mediating the addition of a 3' polynucleotide tail preventing the elongation of the product α' on the aT. This mechanism provides to the system an amplification threshold, meaning that the amplification reaction can start only if a certain concentration of α strands is reached. Below this concentration, the pT acts as a sink that catalytically degrades α strands. Above this threshold concentration, this sink effect is saturated, and the exponential amplification of α starts. The reporting probe (rT) reacts stoichiometrically with α strands to provide a fluorescent signal. All produced strands (α-trigger, α, α') are dynamically degraded by the exonuclease ttRecJ to keep the system out-of-equilibrium (notably to avoid the poisoning of the pT by the accumulation of α').

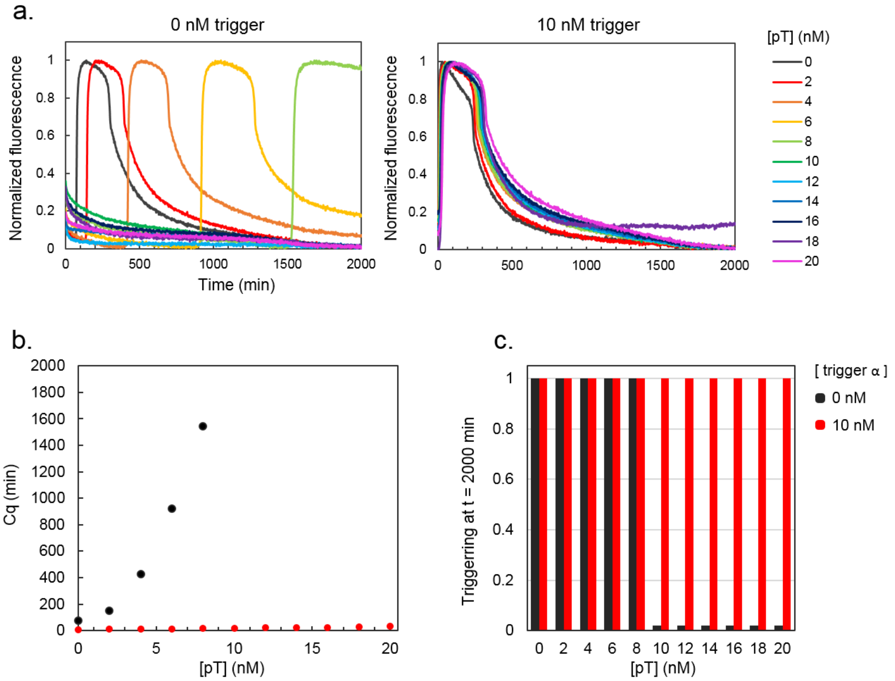


**Supplementary Figure 3. Bistable amplification switch.** The autocatalytic template αtoα mixed with an increasing concentration of pTα is spiked with 0 or 10 nM of trigger α and the amplification is monitored in real-time using the double strand specific binding dye EvaGreen. a. Amplification curves over 2000 minutes. b. The Cq (amplification time, corresponding to the time to reach 20 % of the maximum fluorescence) is plotted as a function of the pseudotemplate concentration. c. Final state of the bistable switch as a function of the initial pT concentration (0 = no amplification, 1 = amplification).

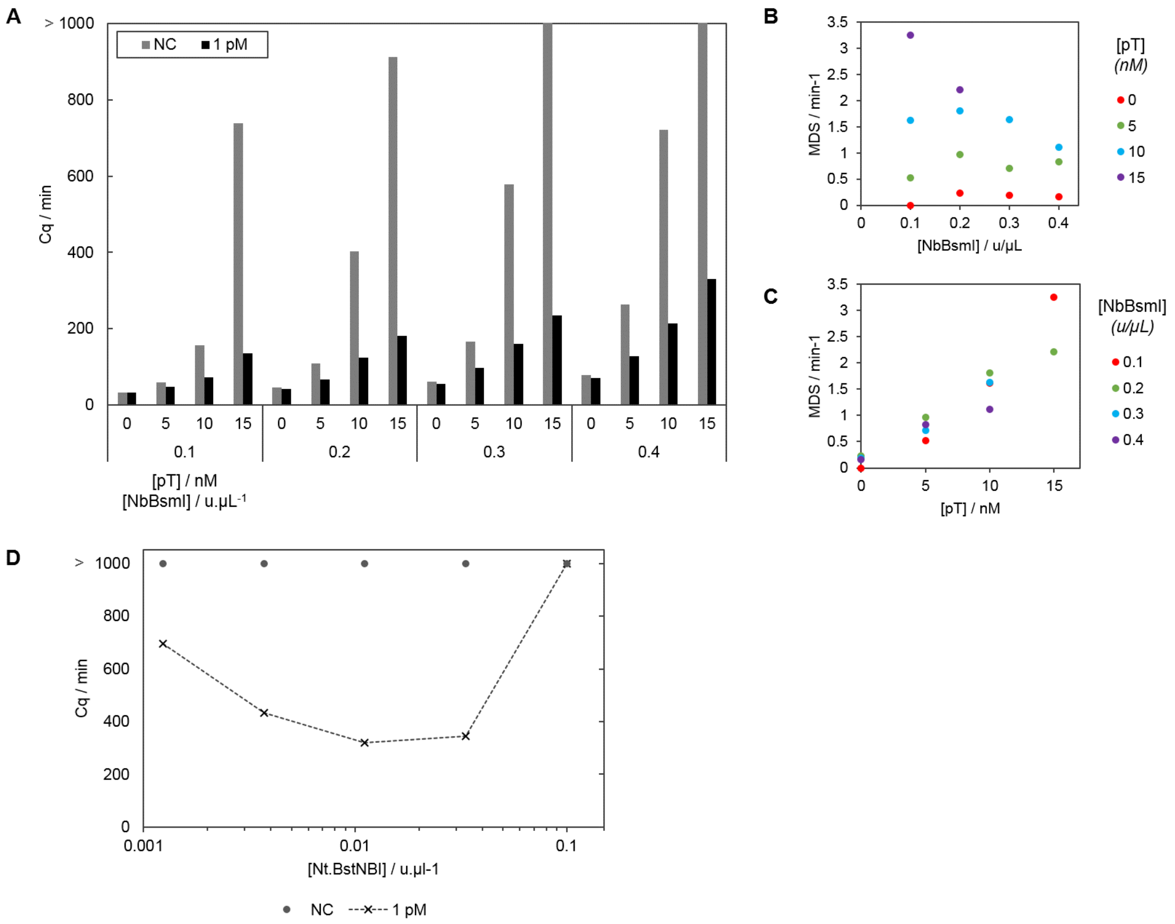


**Supplementary Figure 4. Experimental conditions optimizations.** (**A**) Effect of the pseudotemplate and Nb.BsmI concentrations on the detection of Let-7a. Samples containing a defined concentration of pT (from 0 to 15 nM) and Nb.BsmI (from 0.1 to 0.4 u/µL) are spiked with 0 or 1 pM Let-7a and the amplification reaction was monitored in real-time. The Cq are plotted as a function of the pT and Nb.BsmI concentrations. (**B**) A microRNA detection score (MDS) is calculated for each set of concentration according to the following equation. $\mathrm{MDS} = 100(\frac{\mathrm{Cq_{NC}}}{\mathrm{Cq_{1\,pM}}} - 1)/\mathrm{Cq_{1pM}}$. The MDS is meant to reflect both the time window between the amplification times of the NC and 1 pM and the speed of the target-triggered reaction. The MDS is plotted as a function of b) the pseudotemplate concentration and c) the Nb.BsmI concentration. It is shown that both parameters delay the amplification reaction: the pseudotemplate plays the role of an active sink that degrades part of the produced triggers. Nb.BsmI is known to be inhibited by its own product (the nicked duplex), which probably slows down the reaction by preventing the release of the trigger after cutting the duplex. It is to note that increasing the pseudotemplate concentration positively affect the detection score, by delaying preferentially the NC amplification and affected to a lesser extent the target-triggered amplification. On the other hand, Nb.BsmI affect the amplification time, irrespective from the target concentration and thus negatively affect the detection score. This observation demonstrates the importance of the active degradation mechanism provided by the pseudotemplate to reduce/abolish background amplification. (**C**) Optimization of the Nt.BstNBI concentration. Samples spiked with 0 or 1 pM of Let-7a are incubated in presence of a varying concentration of Nt.BstNBI. The Cq are plotted as a function of the Nt.BstNBI concentration. The graph suggests an optimal concentration around 0.01 u/µL.

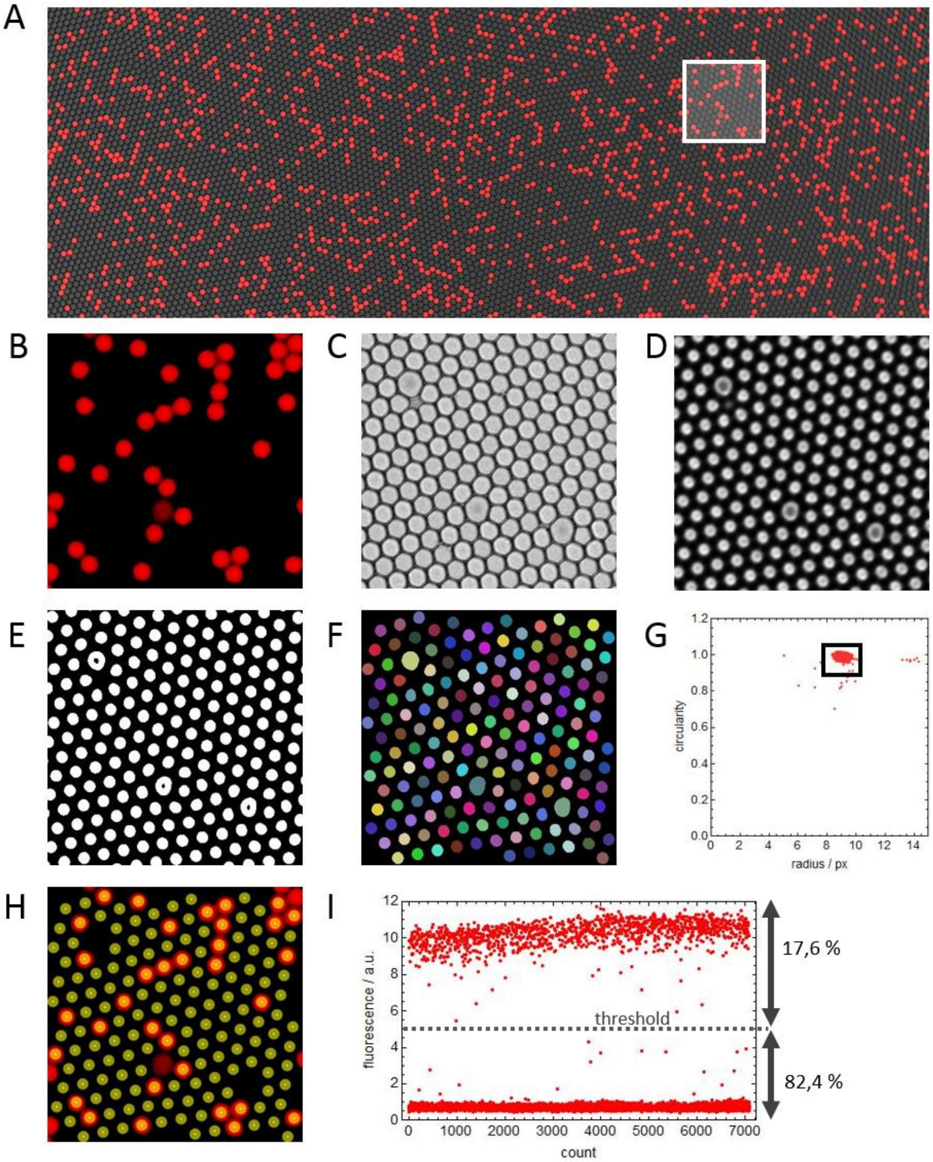


**Supplementary Figure 5: Droplets analysis.** (**A**) After incubation, the emulsion is sandwiched between two hydrophobic glass slides to obtain a monolayer of droplets and imaged with an epifluorescence microscope. (**B**) Atto633 fluorescence channel. (**C**) bright field channel, (**D**) brightfield channel defocused by a 10 µm offset. With this setting, the droplet outlines are reinforced, facilitating the droplets segmentation. (**E**) The brightfield image (D) is binarized and (**F**) all droplets are segmented. (**G**) The droplets are filtered according to their size and circularity. (**H**) The fluorescence of each droplet is extracted from a disk of radius r (with 3 < r < 6 pixels) and center xy (xy corresponding to the centroid of the selected component). (**I**) The positive droplets correspond to the ones with a fluorescence exceeding a set threshold (corresponding to $\frac{\overline{\mathrm{pos}}-\overline{\mathrm{neg}}}{2}$, where $\overline{\mathrm{pos}}$ and $\overline{\mathrm{neg}}$ correspond to the median fluorescence of the positive and negative droplet population respectively). Knowing the droplet volume, the concentration is calculated back from the Poisson law using the equation $[\mathrm{miR}] = \frac{\lambda}{\mathrm{N_A.V}}$, where $\lambda$ represents the average number of microRNA target per droplet, $\mathrm{N_A}$ is the Avogadro number and $\mathrm{V}$ is the volume of the droplets (in L). $\lambda = -\ln(1 - \mathrm{F_{pos}})$, where $\mathrm{F_{pos}}$ is the frequency of positive droplets. The 95% confidence interval is given by the uncertainty on a binomial proportion $\lambda_c = \lambda \pm 1.96\sqrt{\frac{\mathrm{F_{pos}.F_{neg}}}{\mathrm{n_{drop}}}}$, where $\mathrm{F_{pos}}$and $\mathrm{F_{neg}}$ correspond to the frequency of positive and negative droplets respectively and $\mathrm{n_{drop}}$ is the total number of droplets analyzed.

**A**

| Experiment | Expected conc.(fM) | # droplets | # droplets pos | % droplet pos | droplet volume (fL) | λ | | | Measured conc.(fM) | Measured conc. (LOD substracted) |
|---|---|---|---|---|---|---|---|---|---|---|
| XP#1 | 0 | 17026 | 48 | 0.3% | 250 | 2.8E-03 | +/- | 8.0E-04 | 19 | 0 |
| | 8 | 13036 | 60 | 0.5% | 250 | 4.6E-03 | +/- | 1.2E-03 | 31 | 12 |
| | 40 | 4314 | 48 | 1.1% | 250 | 1.1E-02 | +/- | 3.2E-03 | 74 | 55 |
| | 200 | 7293 | 256 | 3.5% | 250 | 3.6E-02 | +/- | 4.9E-03 | 237 | 218 |
| | 1000 | 4833 | 656 | 13.6% | 250 | 1.5E-01 | +/- | 2.0E-02 | 968 | 949 |
| | 5000 | 7831 | 3715 | 47.4% | 250 | 6.4E-01 | +/- | 1.8E-01 | 4267 | 4248 |
| XP#2 | 0 | 12450 | 27 | 0.2% | 200 | 2.2E-03 | +/- | 8.2E-04 | 18 | 0 |
| | 30 | 9559 | 53 | 0.6% | 200 | 5.6E-03 | +/- | 1.5E-03 | 46 | 28 |
| | 125 | 11907 | 223 | 1.9% | 200 | 1.9E-02 | +/- | 2.6E-03 | 157 | 139 |
| | 500 | 16409 | 930 | 5.7% | 200 | 5.8E-02 | +/- | 5.2E-03 | 484 | 466 |
| | 2000 | 20534 | 4969 | 24.2% | 200 | 2.8E-01 | +/- | 4.1E-02 | 2297 | 2279 |
| | 10000 | 24037 | 15175 | 63.1% | 200 | 1.0E+00 | +/- | 3.7E-01 | 8274 | 8256 |
| XP#3 | 0 | 7409 | 11 | 0.1% | 700 | 1.5E-03 | +/- | 8.8E-04 | 4 | 0 |
| | 2.7 | 5871 | 17 | 0.3% | 700 | 2.9E-03 | +/- | 1.4E-03 | 7 | 3 |
| | 11 | 7246 | 53 | 0.7% | 700 | 7.3E-03 | +/- | 2.0E-03 | 17 | 14 |
| | 44 | 7396 | 166 | 2.2% | 700 | 2.3E-02 | +/- | 3.6E-03 | 54 | 50 |
| | 175 | 9258 | 679 | 7.3% | 700 | 7.6E-02 | +/- | 8.1E-03 | 180 | 177 |
| | 700 | 10347 | 2738 | 26.5% | 700 | 3.1E-01 | +/- | 5.1E-02 | 728 | 725 |

**B**

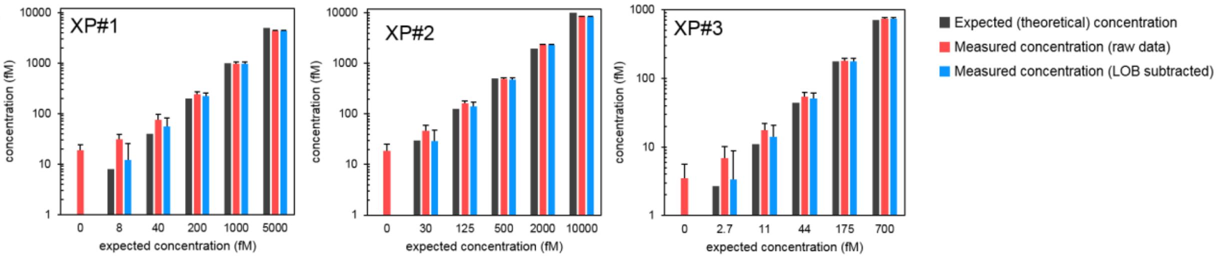


**Supplementary Figure 6: Extended data from the Figure 3d.** (**A**) Raw data obtained from the images analysis for the three independent experiment (XP#1-3). (**B**) the concentration measured is plotted as a function of the expected spiked-in concentration. The limit of the blank (LOB) corresponds to the measured concentration in the negative control. The data are corrected by subtracting the LOB form the measured concentration obtained from the Poisson statistics (cf. Supplementary Figure 5).

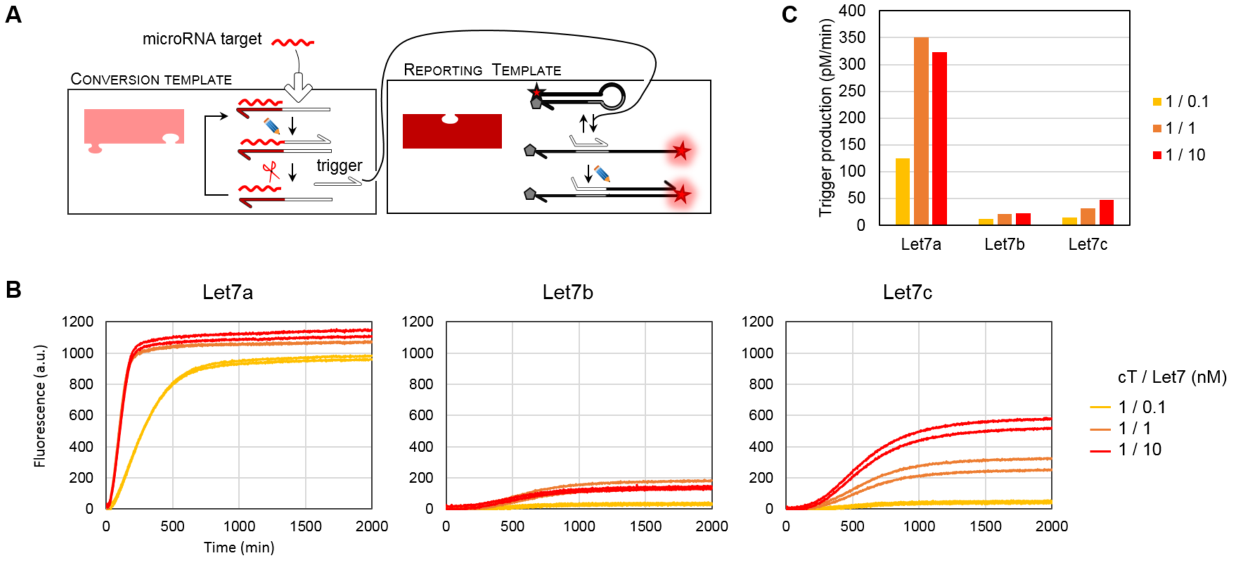


**Supplementary Figure 7. Specificity of the trigger production.** (**A**) The cT Let7atoα (1 nM) is incubated with the target Let7a or analog target Let7b or Let7c (0.1, 1, 10 nM) together with the rTα (50 nM), the polymerase Vent(exo-) and the nicking enzyme NtBstNBI. (**B**) The production of α by the duplexes cT/Let7 is directly visualized by real-time fluorescence monitoring at 50°C. (**C**) The production rate for each condition is estimated as the maximum fluorescence increase of the rTα.

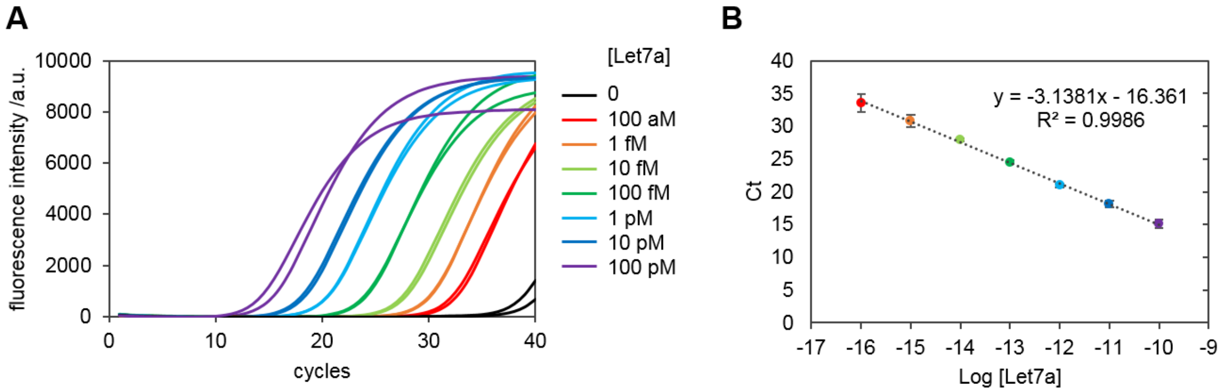


**Supplementary Figure 8. RT-qPCR calibration.** (**A**) Amplification curves using standard concentration of a synthetic Let-7a target obtained for a replicate experiment. (**B**) Ct (cycling threshold) are plotted as a function of the Let-7a concentration.

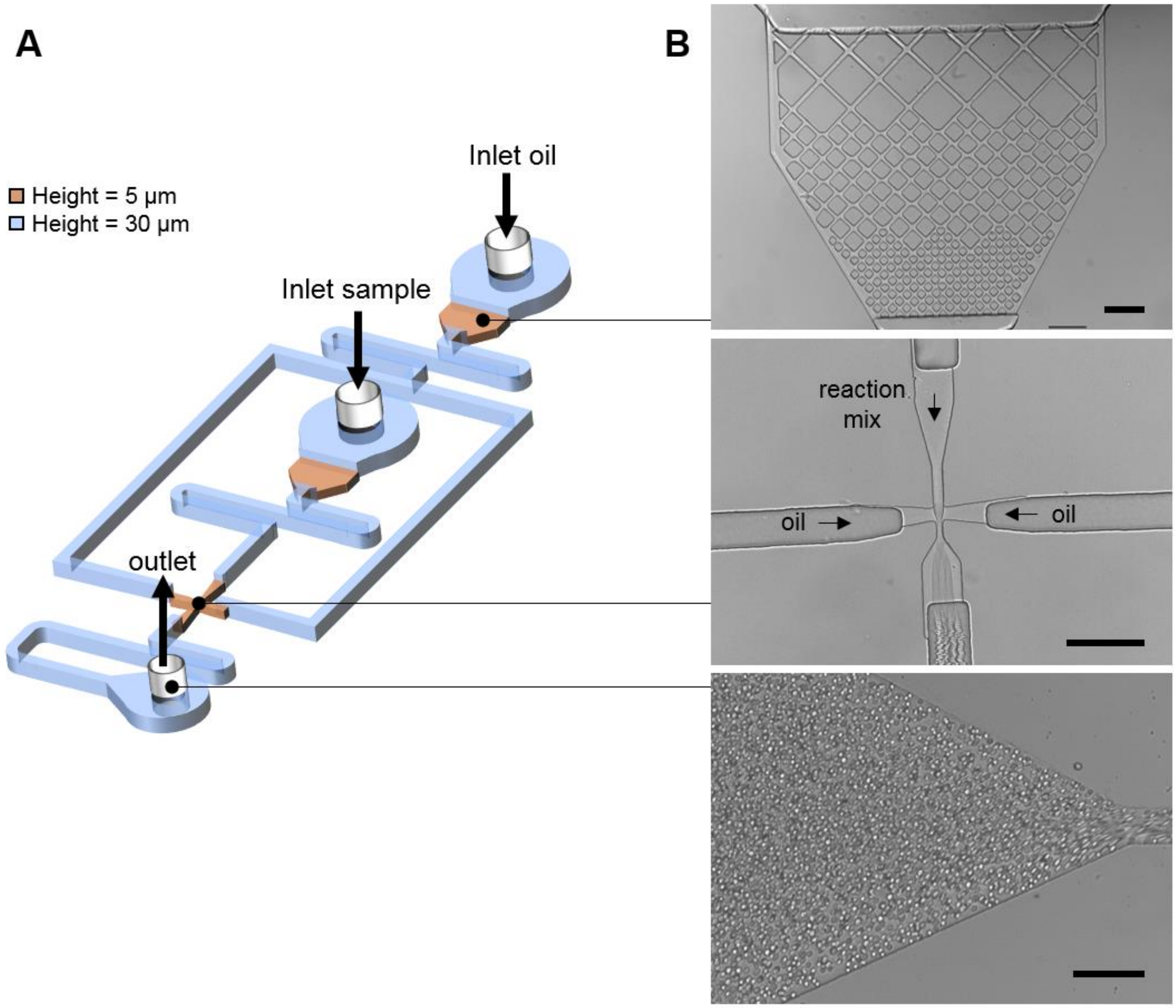


**Supplementary Figure 9. Microfluidic chip design.** (**A**) Layout of the microfluidic circuit. (**B**) Microscopy images of (from top to down) the filter, the flow focusing junction and the outlet. Scale bars represents 100 µm

**Supplementary Table 1. Oligonucleotides sequences used throughout the study**."*" denotes phosphorothioate backbone modification. "p" denotes 3'-phosphate modification. Upper and lower cases represent 2'-deoxyribonucleotide and ribonucleotide, respectively.

| ID | Sequence | Function |
|---|---|---|
| α | CATTCTGGACTG | s-strand |
| αtoα | C*A*G*T*CCAGAATGCAGTCCAGAA p | aT |
| pTα | T*T*T*T*TCAGTCCAGAATG p | pT |
| rTα | Atto633 *A*T*TCTGAATGCAGTCCAGAAT BHQ2 | rT |
| Let7atoα | TGCAGTCCAGAAGTTTGACTCAAACTATACAACCTACTACCTCA p | cT |
| Let7ctoα | TGCAGTCCAGAAGTTTGACTCAAACCATACAACCTACTACCTCA p | cT |
| mir39toα | TGCAGTCCAGAAGTTTGACTCACAAGCTGATTTACACCC p | cT |
| mir92atoα | TGCAGTCCAGAAGTTTGACTCAAGCATTGCAACCGATCCCAACC p | cT |
| mir31*toα | TGCAGTCCAGAAGTTTGACTCAGATGGCAATATGTTGGCA p | cT |
| mir31toα | TGCAGTCCAGAAGTTTGACTCAACAGCTATGCCAGCATCT p | cT |
| mir215toα | TGCAGTCCAGAAGTTTGACTCAGTCTGTCAATTCATAGGTCAT p | cT |
| Let-7a | ugagguaguagguuguauaguu | microRNA |
| Let-7b | ugagguaguagguugugugguu | microRNA |
| Let-7c | ugagguaguagguuguaugguu | microRNA |
| mir39-ce | ucaccggguguaaaucagcuug | microRNA |
| mir92a | agguugggaucgguugcaaugcu | microRNA |
| mir31* | ugcuaugccaacauauugccauc | microRNA |
| mir31 | aggcaagaugcuggcauagcugu | microRNA |
| mir215 | augaccuaugaauugacagac | microRNA |
| Expar-Let7a | AACTATACAACCTACTACCTCAAACAGACTCAAACTATACAACC TACTACCTCAA | EXPAR template |
